## Supplemental Figures for "Non-cell-autonomous control of gastruloid development by the lncRNA *T-UCstem1* through DKK1-dependent modulation of WNT signalling"

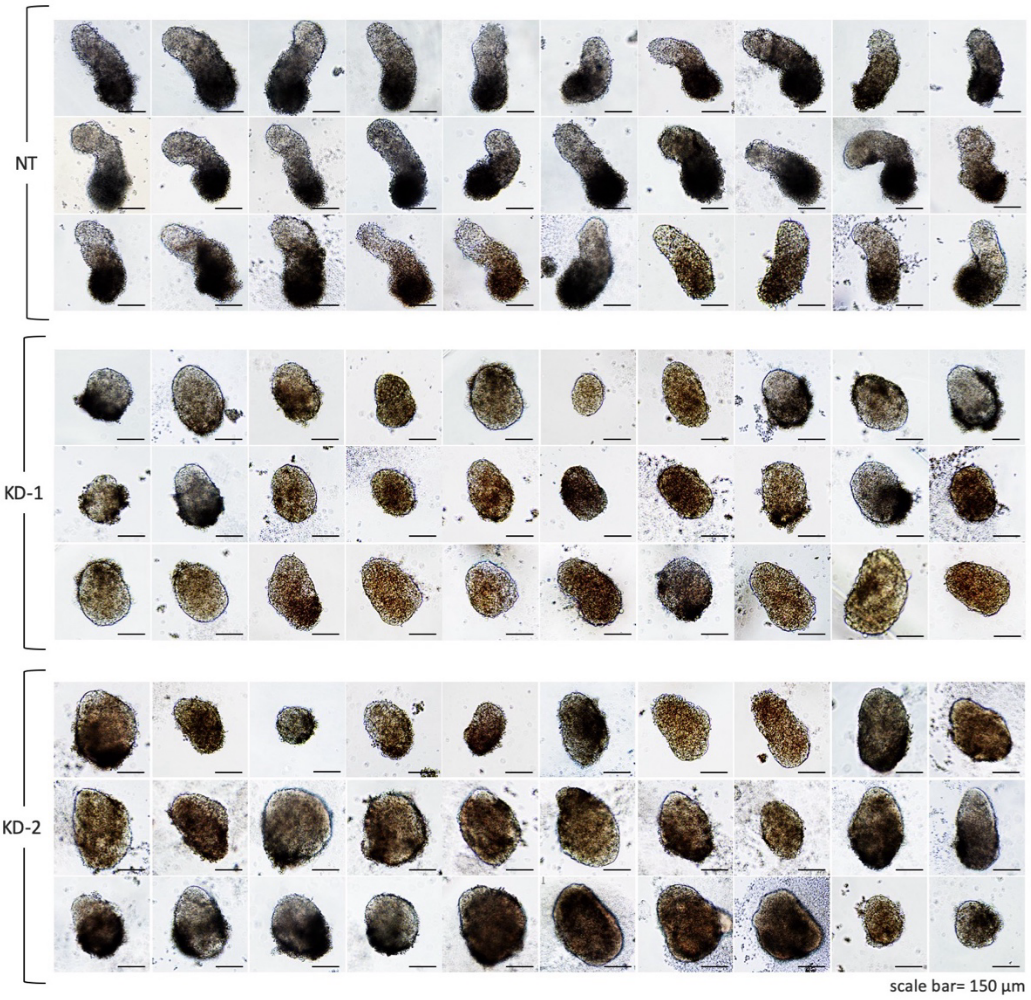


**Figure S1: *T-UCstem1* KD aggregates show a high degree of shape heterogeneity.** The figure showing the heterogeneity of *T-UCstem1* KD gastruloids compared to Control (NT). Scale bar, 150 µm (n=4; 30 gastruloids/condition).

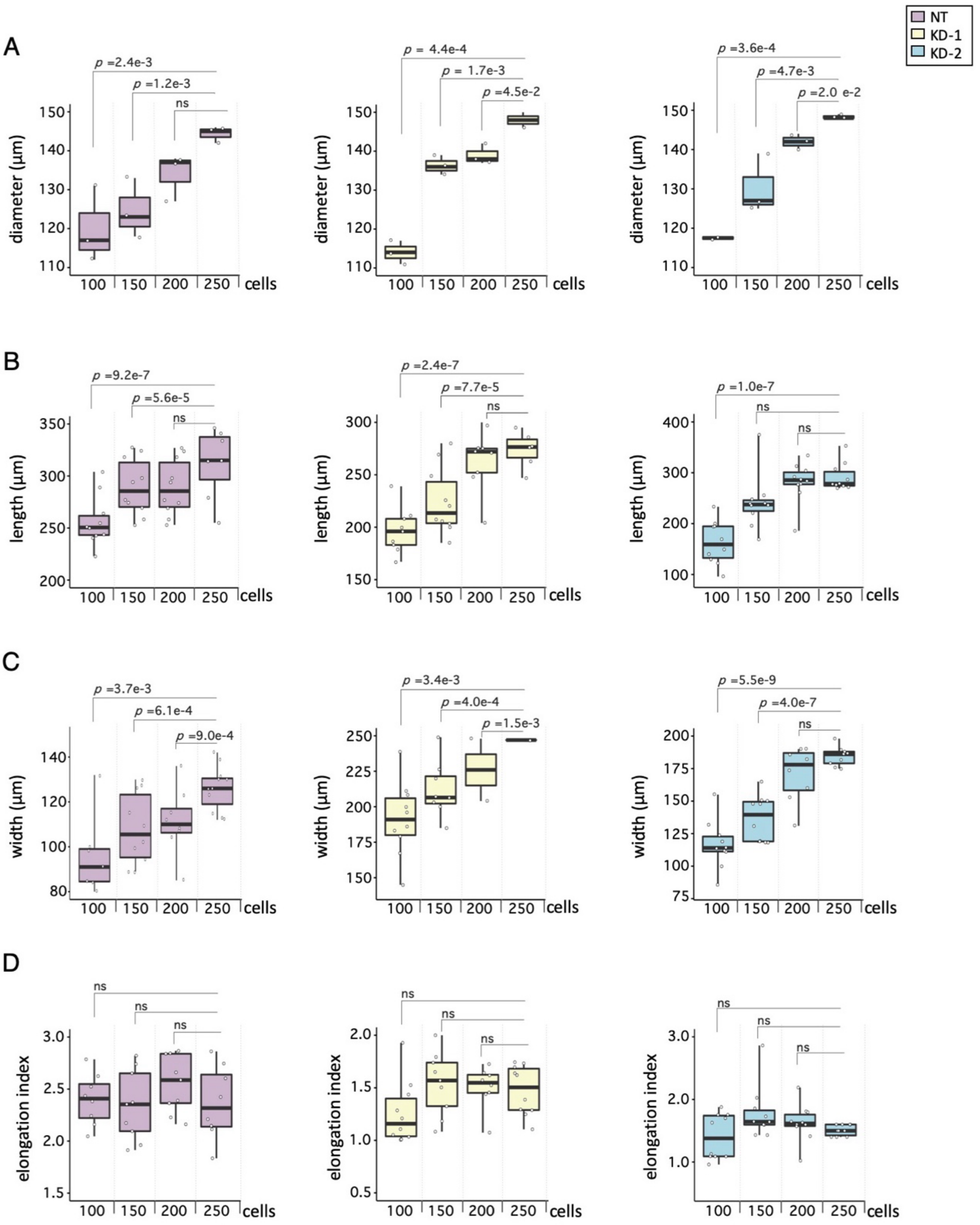


**Figure S2: *T-UCstem1* KD effect is independent on the initial cell number.** A) Boxplot diagrams of the aggregate diameter at 48 h of NT (left), KD-1 (middle) and KD-2 (right). B-D) Boxplot diagrams of the gastruloids length (B), width (C) and elongation index (D) at 120 h of NT (left), *T-UCstem1* KD-1 (middle) and *T-UCstem1* KD-2 (right). Data are shown as the mean ± SD.

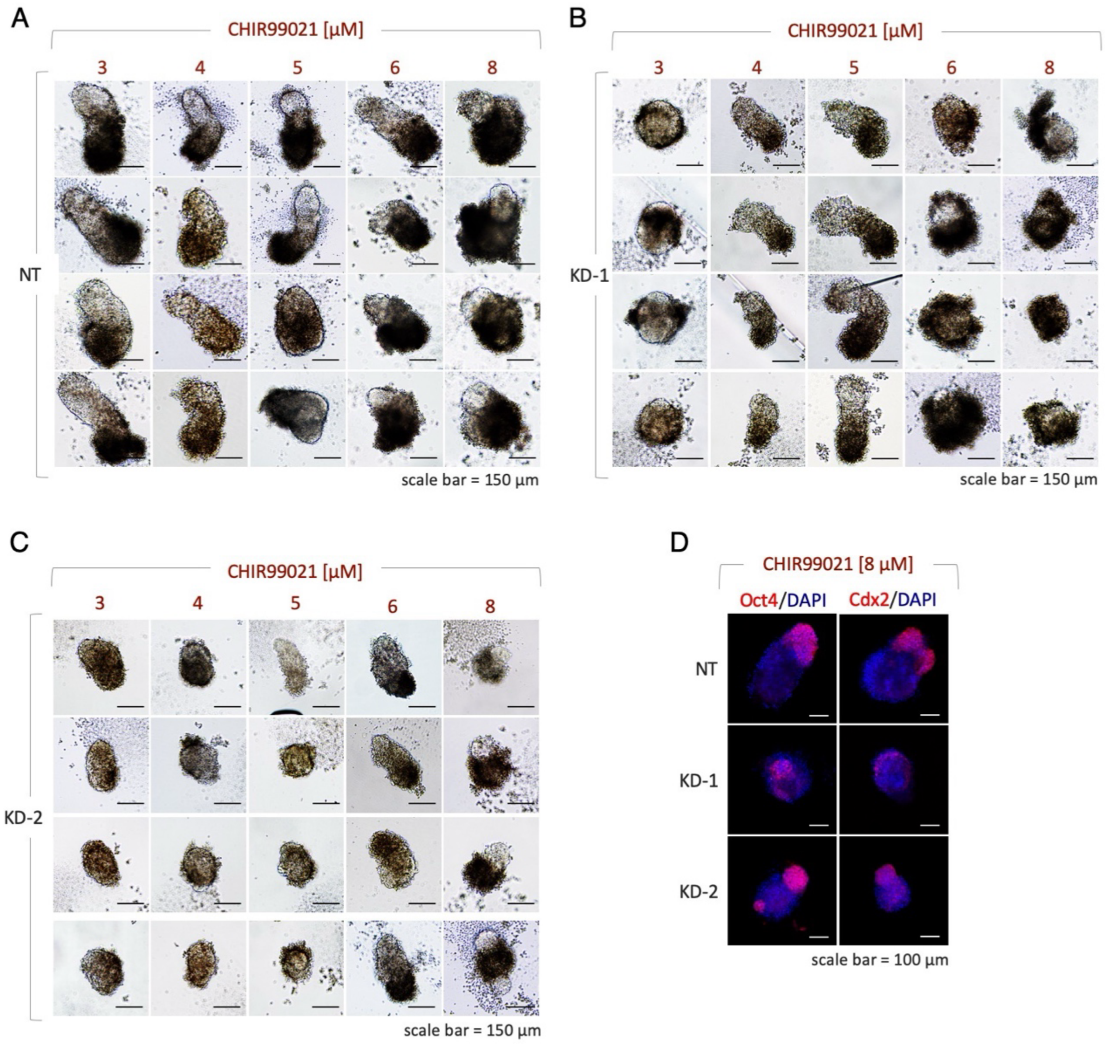


**Figure S3: Dose-dependent effect of CHIR99021 on gastruloid development.** A-C) Representative brightfield images of NT (A), KD-1 (B), KD-2 (C) gastruloids treated with increasing concentration of CHIR99021 (3-8 µM). Scale bar, 150 µm. D) Representative confocal images of Oct4 (pluripotency marker) and Cdx2 (differentiation marker) in gastruloids treated with 8 µM of CHIR99021. Nuclei were counterstained with DAPI (blue). Scale bar, 100 µm.

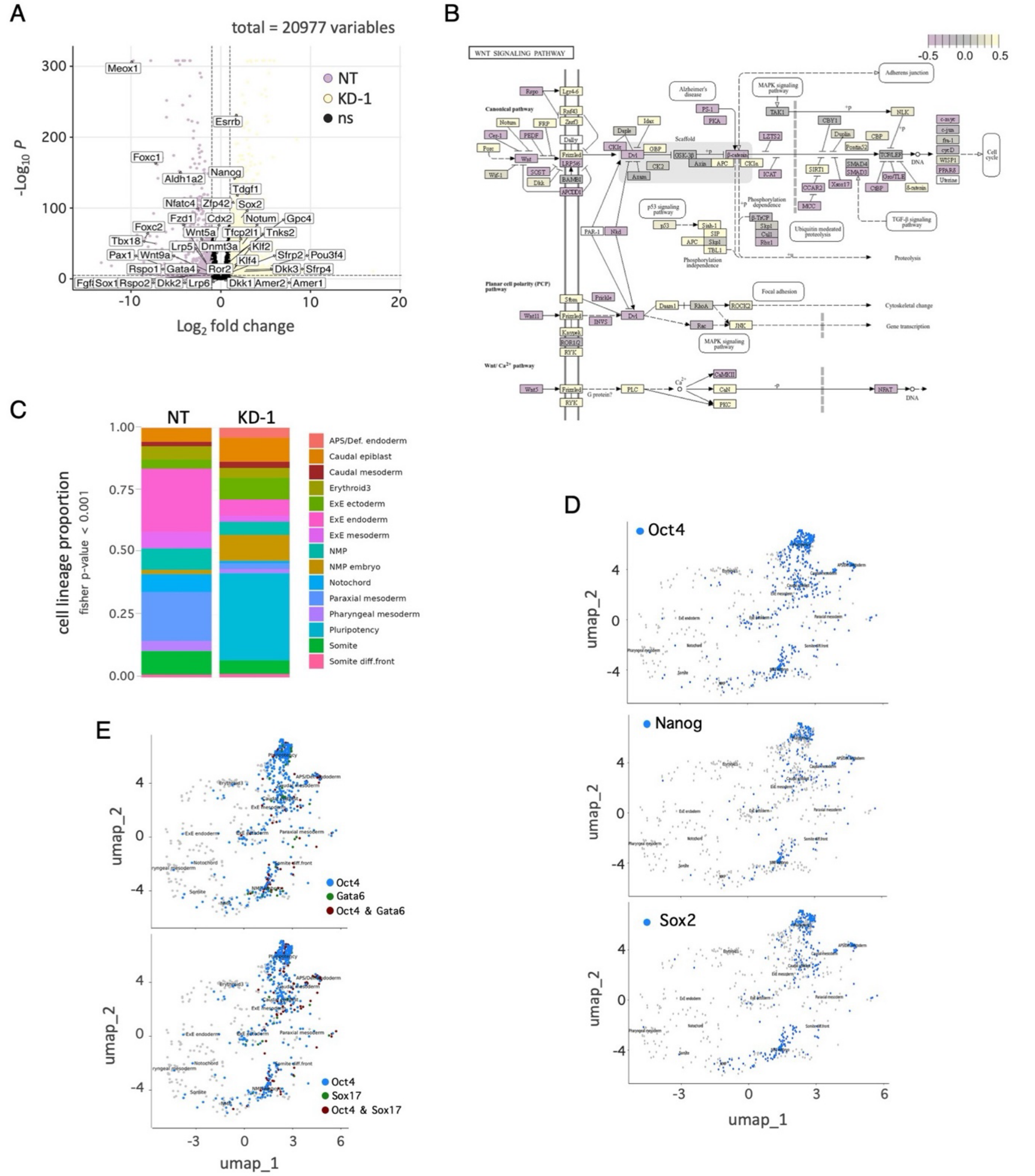


**Figure S4**: **Diverse cellular composition in *T-UCstem1* KD gastruloids versus the Control.** A) Volcano plot of differentially expressed genes between *T-UCstem1* KD-1 and NT gastruloids. B) Reference map of the WNT Signaling pathway from the KEGG pathway database; annotated features are coloured based on differential expression between KD and Ctrl. C) Stacked barplots of the cell lineage proportion in *T-UCstem1* KD-1 and NT gastruloids. D) UMAP plot highlighting expression of *Oct4*, *Nanog* and *Sox2* in *T-UCstem1* KD-1 gastruloids. E) UMAP plot highlighting expression of *Oct4*, *Gata6* and *Sox17* in *T-UCstem1* KD-1 gastruloids.
